# Extreme genetic structure and dynamic range evolution in a montane passerine bird: implications for tropical diversification

**DOI:** 10.1101/376681

**Authors:** Carlos Daniel Cadena, Jorge L. Pérez-Emán, Andrés M. Cuervo, Laura N. Céspedes, Kevin L. Epperly, John T. Klicka

**Affiliations:** Laboratorio de Biología Evolutiva de Vertebrados, Departamento de Ciencias Biológicas, Universidad de Los Andes, Bogotá, Colombia; Instituto de Zoología y Ecología Tropical, Universidad Central de Venezuela, Caracas, Venezuela; Colección Ornitológica Phelps, Caracas, Venezuela; Louisiana State University Museum of Natural Science; Instituto de Investigación en Recursos Biológicos Alexander von Humboldt; Department of Biology and Burke Museum of Natural History and Culture, University of Washington, Seattle

**Keywords:** Andes, elevational replacement, range expansion, speciation

## Abstract

**Aim:** Employ phylogeographic analyses of a widespread species complex to examine the role of historical and evolutionary processes in the origin and maintenance of high species diversity in the Neotropical montane region.

**Location:** Neotropical highlands.

**Taxon:** *Henicorhina* wood-wrens (Aves, Troglodytidae).

**Methods:** We collected mtDNA sequence data for 288 individuals thoroughly covering the range of the *Henicorhina leucophrys* complex from Mexico to Bolivia. Sequences were employed to characterize population structure, infer phylogenetic relationships among populations and their divergence times, examine lineage accumulation through time, and identify presumptive species using coalescent methods. We also explored the origin of elevational and latitudinal replacements involved in spatial changes in species assemblages in the Andes.

**Results:** We found remarkable genetic structure within the complex, which consists of numerous lineages reaching >12% sequence divergence; most divergent populations occur in areas separated by topographic barriers but several of them, typically not sister to each other, co-occur with elevational segregation on mountain slopes or replace each other with latitude along the Andes. Some close relatives occur in areas separated by thousands of kilometers, with more distant relatives occupying intervening areas. The complex likely originated in the Mexican highlands and expanded extensively in South America while diverging rapidly at a constant rate into many different lineages which have persisted for millions of years. Coalescent analyses consistently revealed that the complex may comprise more than 30 species; while we do not suggest these presumptive species should be recognized by taxonomists in the absence of additional data, *H. leucophrys* is a distant outlier among New World birds in terms of high lineage diversity within a single recognized species.

**Main Conclusions:** Our study captured wood-wren lineages in the act of building up diversity via divergence and persistence in allopatry, achievement of secondary sympatry, and coexistence at the landscape scale mediated by ecological and evolutionary divergence. Although dispersal by wood-wrens is restricted at present and this likely accounts for strong population structure across topographic barriers, their ranges have been dynamic, managing to disperse over much of the montane Neotropics. Phases of expansion and contraction of ranges and localized extinctions of populations likely account for phylogeographic patterns which are precursors to the origin of new species and the accumulation of diversity in tropical mountains.

## Introduction

Mountains in the tropics contribute disproportionately to regional species richness given their area in comparison to lowlands, and are often considered global hotspots of biological diversity and endemism (Stattersfield *et al*., 1997; Orme *et al*., 2005; Fjeldså *et al*., 2012). Tropical mountains exhibit particularly high beta diversity (i.e. species turnover in space) because (1) species assemblages shift along elevational gradients, and (2) related species occupy similar elevations in different mountains or in sectors of a mountain separated by geographic barriers. While knowledge of the biodiversity of mountains has advanced conceptually (Graham *et al*., 2014; Bertuzzo *et al*., 2016; Badgley *et al*., 2017) and empirically (e.g., Patterson *et al*., 1998; Jankowski *et al*., 2009; McCain, 2009; Price *et al*.,2014; Peters *et al*., 2016; Quintero & Jetz, 2018), accounting for species richness in montane systems remains difficult. In particular, although climate and available energy have an imprint globally on the distribution of life (Francis & Currie, 2003; Hawkins *et al*., 2003), they cannot predict the agglomeration of range-restricted species in tropical mountains (Rahbek & Graves, 2001; Jetz & Rahbek, 2002; Rahbek *et al*., 2007; but see Ruggiero & Hawkins, 2008). Because the high richness and uniqueness of mountains in the tropics may instead reflect high speciation rates or low extinction rates (Jetz *et al*., 2004; Badgley *et al*., 2017), considering evolutionary processes is crucial to a better understanding of montane diversity (Graham *et al*., 2014; Laiolo *et al*., 2018; Quintero & Jetz, 2018).

Dozens of studies have used phylogenetic and population genetic perspectives to probe into evolutionary processes underlying patterns of avian diversity in the Neotropical mountains. Birds have diversified rapidly in the Andes, with pivotal roles of features of the landscape (e.g., low-lying valleys, high-elevation passes) and of climatic changes as drivers of divergence (Pérez-Emán, 2005; Weir, 2006; Cadena *et al*., 2007; Ribas *et al*., 2007; Sedano & Burns, 2010; Chaves *et al*., 2011; Gutiérrez-Pinto *et al*., 2012; Valderrama *et al*., 2014; Benham *et al*., 2015; Sánchez-González *et al*., 2015; Winger & Bates, 2015; Prieto-Torres *et al*., 2018). Allopatric differentiation of lineages separated by barriers to dispersal is predominant (Hazzi *et al*., 2018), whereas evidence for speciation in parapatry along mountain slopes remains elusive (Patton & Smith, 1992; García-Moreno & Fjeldså, 2000; Cadena *et al*., 2012; Caro *et al*., 2013). Thus, the replacement of closely related species along elevational gradients, a salient geographic pattern in tropical avifaunas (Terborgh, 1971, 1977), appears to result largely from populations coming into secondary contact after allopatric divergence (Diamond, 1973; Cadena, 2007; Freeman, 2015). However, with hundreds of bird species living in the Neotropical mountains, much remains to be learned about the histories of individual clades and about how such histories collectively resulted in the patterns of diversity we observe today.

Before conducting analyses seeking to characterize and account for patterns of diversity one must have proper knowledge of what species exist and where they occur (Fine, 2015). Traditionally, the species-level taxonomy of birds was considered well-known (Scheffers *et al*., 2012), with suggestions that the inventory of species was essentially complete by the mid 20th century (Mayr, 1946). This, however, proved incorrect: multiple avian species have been discovered and described over recent decades, and analyses of novel data sets (notably, of vocal and genetic variation) have revealed that species-level diversity was seriously underestimated (Fjeldså, 2013). The extent to which avian taxonomy will require revision depends on how one delimits species (Tobias *et al*., 2010; Gill, 2014; Toews, 2015; Barrowclough *et al*., 2016; Remsen, 2016), but clearly there are more species of birds than traditionally thought, particularly in the tropics. Although problems with species delimitation are unlikely to affect assessments of patterns in local (alpha) diversity of birds, inadequate knowledge of species limits may seriously influence perceptions of patterns in species turnover in space and hence regional and global patterns of diversity (beta and gamma diversity). Alternative approaches for species delimitation may also influence inferences about biogeographic history (Smith *et al*.,2018).

Birds in which species diversity is likely greater than traditionally thought are those in which plumages vary subtly (in which case one would expect species recognition to be based more on vocal cues), and in which ecologically relevant traits (body size, habitat, dispersal ability) may be conducive to population isolation (Burney & Brumfield, 2009; Salisbury *et al*., 2012; Smith *et al*., 2014; Harvey *et al*., 2017a). Here, we analyze the phylogeography of the Grey-breasted Wood-wren complex *(Henicorhina leucophrys,* Troglodytidae), a group of small, drably colored and highly vocal songbirds of forest interior, with poor dispersal abilities. Because the complex is broadly distributed from Mexico to Bolivia and restricted to montane forest habitats, it is an appropriate system in which to ask questions relevant to understanding the role of evolutionary processes in establishing patterns of diversity in Neotropical mountains. We used extensive geographic sampling to reconstruct the phylogenetic relationships of populations in the complex and to characterize patterns of genetic variation with the goals of (1) gaining insight about the tempo and mode of evolutionary differentiation and on the role of colonization of new regions in diversification, (2) understanding the role of geographic isolation in the differentiation of lineages, and (3) exploring the origin of elevational replacements leading to changes in species assemblages with elevation. We also examined the extent to which current taxonomy adequately reflects true diversity and reflected on the influence of cryptic differentiation for inferences about diversification processes and patterns of diversity in the tropics.

## Methods

### Study system

*Henicorhina* wrens (Troglodytidae) range widely in the Neotropical region. Traditionally, taxonomists recognized two widespread species, the White-breasted Wood-Wren (*H. leucosticta*) and the Grey-breasted Wood-Wren (*H. leucophrys*). Two species with restricted ranges, Bar-winged Wood-Wren (*H. leucoptera*) from southern Ecuador and northern Peru (Fitzpatrick *et al*., 1977), and Munchique Wood-Wren (*H. negreti*) from western Colombia (Salaman *et al*., 2003), were later described. More recently, another narrow endemic formerly considered a subspecies of *H. leucophrys*, Hermit Wood-Wren (*H. anachoreta*) from northern Colombia, was elevated to species status (Cadena *et al*., 2016). Preliminary data on phylogenetics and population structure of wood-wrens based on mitochondrial DNA sequences suggest that both *H. leucosticta* and *H. leucophrys* are paraphyletic (*H. leucoptera* is nested within *H. leucosticta* and *H. anachoreta* is nested within *H. leucophrys)*, and both comprise multiple distinct lineages (Dingle *et al*., 2006; Becker *et al*., 2007; Caro *et al*., 2013; Aguilar *et al*., 2014; Smith *et al*., 2014). However, no comprehensive analysis of genetic variation across the range of either widespread species has been conducted.

Wood-wrens segregate ecologically by elevation. Overall, *H. leucosticta* is a lowland species replaced in montane areas by *H. leucophrys;* their replacement is sharp and likely mediated by interspecific competition (Jankowski *et al*., 2010). In the isolated Cordillera del Cóndor of southern Ecuador and northern Peru, *H. leucophrys* also replaces *H. leucosticta* in montane areas but is absent from higher elevations where *H. leucoptera* occurs (i.e. the three species turn over along the elevation gradient; Fitzpatrick *et al*., 1977; Dingle *et al*., 2006). Likewise, in part of the western slope of the Colombian Andes, *H. negreti* replaces *H. leucophrys* (subspecies *brunneiceps*) at higher elevations, and is in turn replaced by nominate *H. leucophrys* east of the ridgeline on the eastern slope of the cordillera (Salaman *et al*.,2003). In addition, two populations of *H. leucophrys* differing in mtDNA sequences, morphology, and songs are parapatrically distributed along an elevational gradient in Ecuador (Dingle *et al*., 2008; Dingle *et al*., 2010), but nuclear gene flow indicates they are conspecific (Halfwerk *et al*., 2016). A similar scenario with populations differing genetically, morphologically and vocally, and turning over along an elevational gradient exists in the Sierra Nevada de Santa Marta, northern Colombia (Caro *et al*., 2013; Burbidge *et al*., 2015); because there is little to no hybridization, these populations are now treated as separate species, with *H. anachoreta* sharply replacing *H. leucophrys* at higher elevations (Cadena *et al*., 2016).

### Sampling

We focused on the *H. leucophrys* complex, i.e. the clade defined by the most recent common ancestor of populations referable to *H. leucophrys* in current taxonomy and *H. anachoreta* (Dingle *et al*., 2006; Caro *et al*., 2013). Although *H. negreti* was not sampled in previous molecular analyses, we consider it part of the complex based on phenotypic traits (Salaman *et al*., 2003) and our data (see below). The *H. leucophrys* complex is widespread in Neotropical mountains, ranging from Mexico to Bolivia (Figure 1); as currently circumscribed, it consists of 19 taxa including *H. anachoreta, H. negreti*, and 17 subspecies of *H. leucophrys* (Kroodsma & Brewer, 2005). For phylogeographic analyses, we sought to sample as thoroughly as possible across geography and taxonomy. Combining sequences generated for this study and published sequences available in GenBank (total 288 individuals), we managed to cover nearly all of the distribution range of the complex and all named taxa, with multiple individuals and localities per taxon whenever possible (Figure 1; Supplementary Table 1). Sampling in Middle America covered all the major montane areas where members of the complex occur; within South America sampling was especially thorough in Venezuela, Colombia and Ecuador, and less so in the southern part of the range (i.e. Peru and Bolivia). As outgroups for phylogenetic analyses, we used specimens of *H. leucosticta* and *H. leucoptera*, and of species of *Microcerculus, Campylorhynchus*, *Cistothorus, Troglodytes, Cantorchilus* and *Cyphorhinus* (see Barker, 2017 for an overview of relationships among wren genera), for a grand total of 300 individuals considered in analyses.

**Figure 1.**
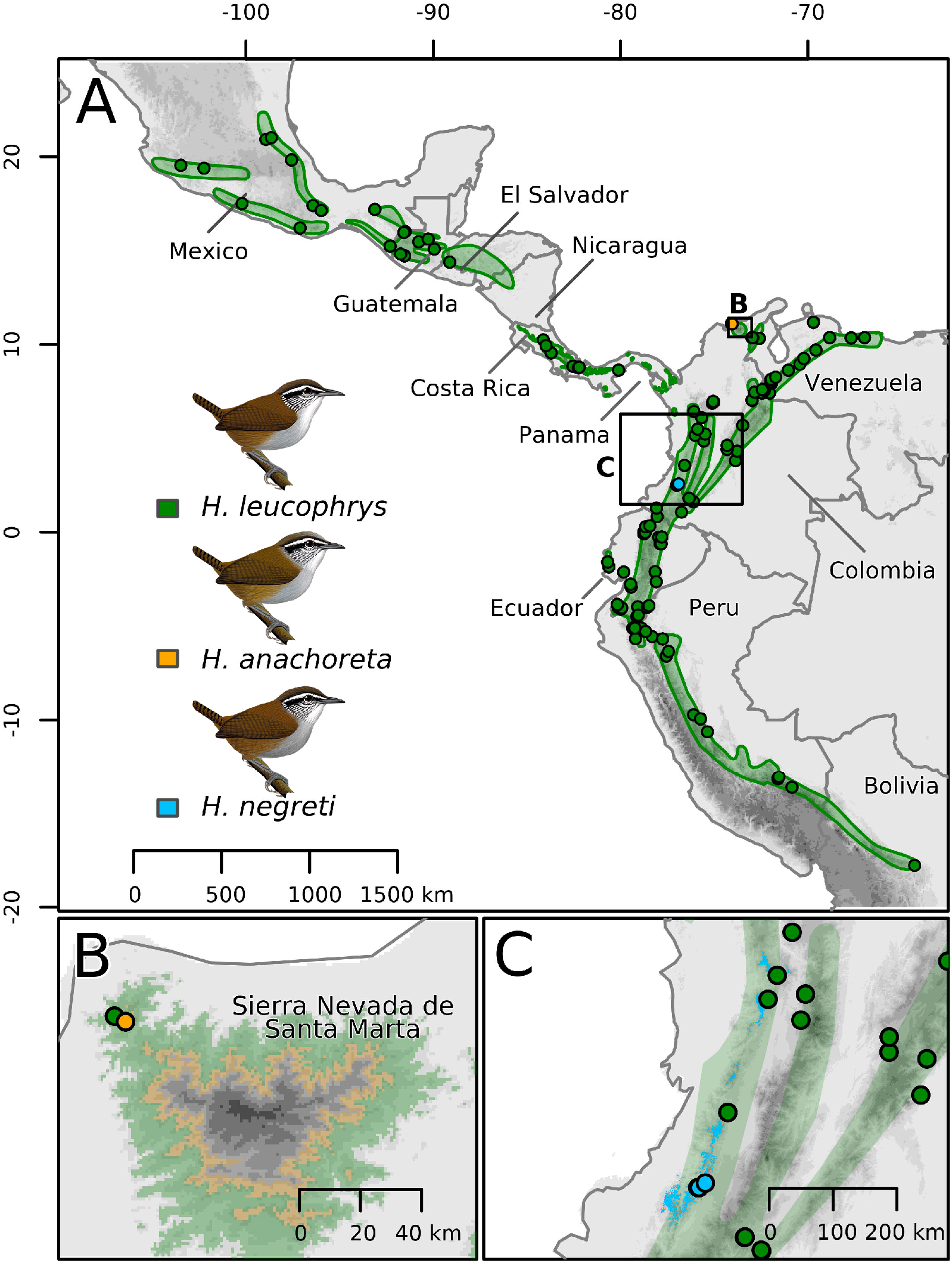
Geographic distribution of wood-wrens in the *Henicorhina leuocophrys* complex in the Neotropical montane region and localities where specimens were sampled for our phylogeographic analyses. The complex currently comprises three species: the widely distributed *H. leucophrys* ranging from Mexico to Bolivia and two narrow endemics from Colombia (*H. anachoreta* in the Sierra Nevada de Santa Marta and *H. negreti* on the western slope of the Cordillera Occidental). Distribution maps were obtained from BirdLife International (*H. leucophrys*) and Velásquez-Tibatá et al. (2013; *H. negreti*), or generated for this study based on information on elevational range (*H. anachoreta*; Cadena *et al*., 2016). Illustrations by F. Ayerbe.

### DNA extraction and sequencing

We extracted DNA using DNeasy tissue extraction kits (Qiagen, Valencia, CA) following the manufacturer’s protocol. We then amplified an 842 base-pair region of the mtDNA gene ATPase 6/8 using primers described by Joseph et al. (2004). We chose to sequence this region because it has been employed in earlier studies of *Henicorhina* (Dingle *et al*.,2006; Dingle *et al*., 2008; Caro *et al*., 2013), which enabled us to include published sequences in analyses. Fragments were amplified via polymerase chain reaction (PCR) in 12.5 μl reactions with denaturation at 94 °C for 10 min, 40 cycles of 94 °C for 30 s, 54 °C for 45 s, and 72 °C for 2 min, followed by 10 min elongation at 72 °C and 4 °C soak. PCR products were sequenced at the Barrick Museum of Natural History (University of Nevada, Las Vegas), Universidad de los Andes (Bogotá, Colombia), or the High-Throughput Genomics Unit at the University of Washington. Chromatograms were aligned using Sequencher v4.9 (GeneCodes Corporation, Ann Arbor, MI).

### Analyses

#### Gene trees

Before phylogenetic analyses, we determined the best-fit model of evolution to be GTR + G with jModeltest v2. 1. 7 (Posada, 2008). We then used Bayesian (BEAST v1.8.4; Drummond *et al*., 2012) and maximum-likelihood (RAxML v 8.2.4; Stamatakis, 2006) methods to estimate phylogenetic trees. For the Bayesian analyses we ran 50 million generations, sampling trees and parameters every 5000 generations. A relaxed uncorrelated lognormal clock with a rate of 2% and a birth-death speciation tree prior (Ritchie *et al*., 2017) were applied. We confirmed likelihood stationarity and adequate effective sample sizes above 200 for all estimated parameters using Tracer v1.6.0 (http://tree.bio.ed.ac.uk/software/tracer). The parameter values of the samples from the posterior distribution on the maximum clade credibility tree were summarized after discarding the first 5 million generations (10%) as burn-in using TreeAnnotator v1.8.4 (Drummond *et al*., 2012). Maximum-likelihood analyses were conducted using a GTRGAMMA model and run for 1000 nonparametric rapid bootstrap replicates to provide an assessment of nodal support.

Both Bayesian and maximum-likelihood analyses were first done with a dataset containing all ATPase sequences (n = 300), then repeated with a reduced data set containing only individuals having unique haplotypes (n = 184), and then reduced further by removing all non-*Henicorhina* taxa (i.e. outgroups). Trees constructed using the latter data set (n = 178) were set aside for use in species delimitation analyses described below. To visualize and annotate trees and to produce figures, we employed R packages ggtree (Guangchuang *et al*., 2017) and phytools (Revell, 2012), and QGIS v 2.18.20 with the Qgis2threejs plugin (http://qgis.osgeo.org).

#### Species delimitation

Given uncertainty about species diversity in the *H. leucophrys* complex, we employed two coalescent approaches using mtDNA data to identify distinct lineages which may be considered presumptive species worthy of additional study with other sources of information (Carstens *et al*., 2013; Sukumaran & Knowles, 2017): (1) multi-rate Poisson Tree Processes (mPTP v0.2.0; Kapli *et al*., 2017) and (2) a Bayesian implementation of the General Mixed Yule Coalescent Model (bGMYC v1.0.2; Reid & Carstens, 2012). The trees generated by BEAST using the unique haplotype dataset were used as input for these analyses.

For the mPTP analysis we ran five replicate mcmc chains of 10,000,000 generations, sampling every 10,000 with a burn-in of 1,000,000 (10%) using the maximum clade credibility tree obtained as described above. Minimum branch length was calculated using the minbr_auto function prior to the analysis. We ran bGMYC with 100 of the 10,000 gene trees estimated in the BEAST analysis, after removing 10% as burn-in; this approach accounts for error in gene-tree estimation by integrating over uncertainty in tree topology and branch lengths. For each tree we ran an MCMC chain of 50,000 steps with 40,000 steps of burn-in and a thinning interval of 100 steps. We focused our results and discussion on lineages (i.e. presumptive species) defined using a threshold of 0.5 on probability of membership of individuals (Gehara *et al*., 2017). However, we also considered more conservative approaches where presumptive species were identified as clusters in the gene tree with posterior probabilities of belonging to the same species ≥0.90 or ≥ 0.95, which allowed us to compare our results to similar work on other birds (Smith *et al*., 2014; Harvey *et al*.,2017b; Smith *et al*., 2017).

#### Diversification through time

To describe patterns of lineage accumulation over time, we constructed lineage-through-time (LTT) plots and estimated the gamma statistic (Pybus *et al*., 2000). We accounted for phylogenetic uncertainty by performing these analyses on a sample of credible trees in the posterior distribution obtained from the BEAST analysis. We employed the 100 trees constructed using only unique haplotypes as input for the bGMYC analyses and trimmed them to include 39 tips corresponding to the presumptive species recognized under the 0.5 threshold. Then we used functions implemented in R packages ape (Paradis *et al*., 2004) and paleotree (Bapst, 2012) to build an LTT plot with a 95% confidence interval and to calculate the gamma statistic for each tree. We also examined diversification dynamics employing Bayesian Analysis of Macroevolutionary Mixtures (program BAMM v 2.5.0; Rabosky *et al*., 2013). Because results were qualitatively similar between methods, we report only those obtained using the simpler approach implemented in ape.

## Results

We found substantial genetic differentiation among populations in the *H. leucophrys* complex. In total, we recovered 172 haplotypes among the 288 individuals analyzed. Several haplotypes were highly divergent from each other, with uncorrected genetic distances between them reaching >12 % (i.e. between individuals from Sierra Madre del Sur of Mexico and from the east slope of the Cordillera Occidental of Colombia). Genetic variation was highly structured spatially, but was not readily accounted for by geographic distance among populations. We did not conduct formal analyses of isolation by distance, but genetic distances among isolated populations from adjacent mountains were often much greater than genetic distances observed over larger distances in more continuous ranges. For instance, mean genetic distances among the five montane areas of Venezuela that we sampled was 7.3% (range 6.1%-8.3%), whereas genetic distances within montane regions extending over comparable distances were much lower, e.g. reaching only 3.5% in the Sierra Madre Oriental of Mexico or 1.7% in the Talamanca-Chiriquí mountains of Costa Rica and Panama.

Maximum-likelihood and Bayesian phylogenetic analyses recovered similar overall patterns (Figure 2., Supplementary Figures 1-2). The deepest split in gene trees separates clades corresponding to Mexican populations from the Sierra Madre del Sur and the western reaches of the Trans-Mexican Volcanic Belt from a large and strongly supported clade including the remainder of populations in the complex. Within the latter clade, the earliest diverging group occurs in eastern Mexico and consists of three subclades, each corresponding to a unique region within the Sierra Madre Oriental. Sister to this group is a clade divided in two main groups (albeit without strong support, i.e. 0.88 posterior probability, 68% ML bootstrap in analyses using only unique haplotypes): one includes samples from lower Central America (Costa Rica and Panama, subspecies *collina)*, whereas the other includes all South American populations of *H. leucophrys*, the Colombian endemic species *H. anachoreta* and *H. negreti*, and a clade formed by samples of *H. leucophrys* from Chiapas (Mexico), Guatemala, and El Salvador (subspecies *castanea* and *composita*). The latter clade, of somewhat uncertain affinities within an otherwise South American group (it was recovered as sister to *H. negreti* with 0.95 posterior probability and 56% ML bootstrap in analyses using only unique haplotypes), is the only exception to the pattern in which Mexican and Central American populations are the earliest diverging lineages in the complex.

**Figure 2.**
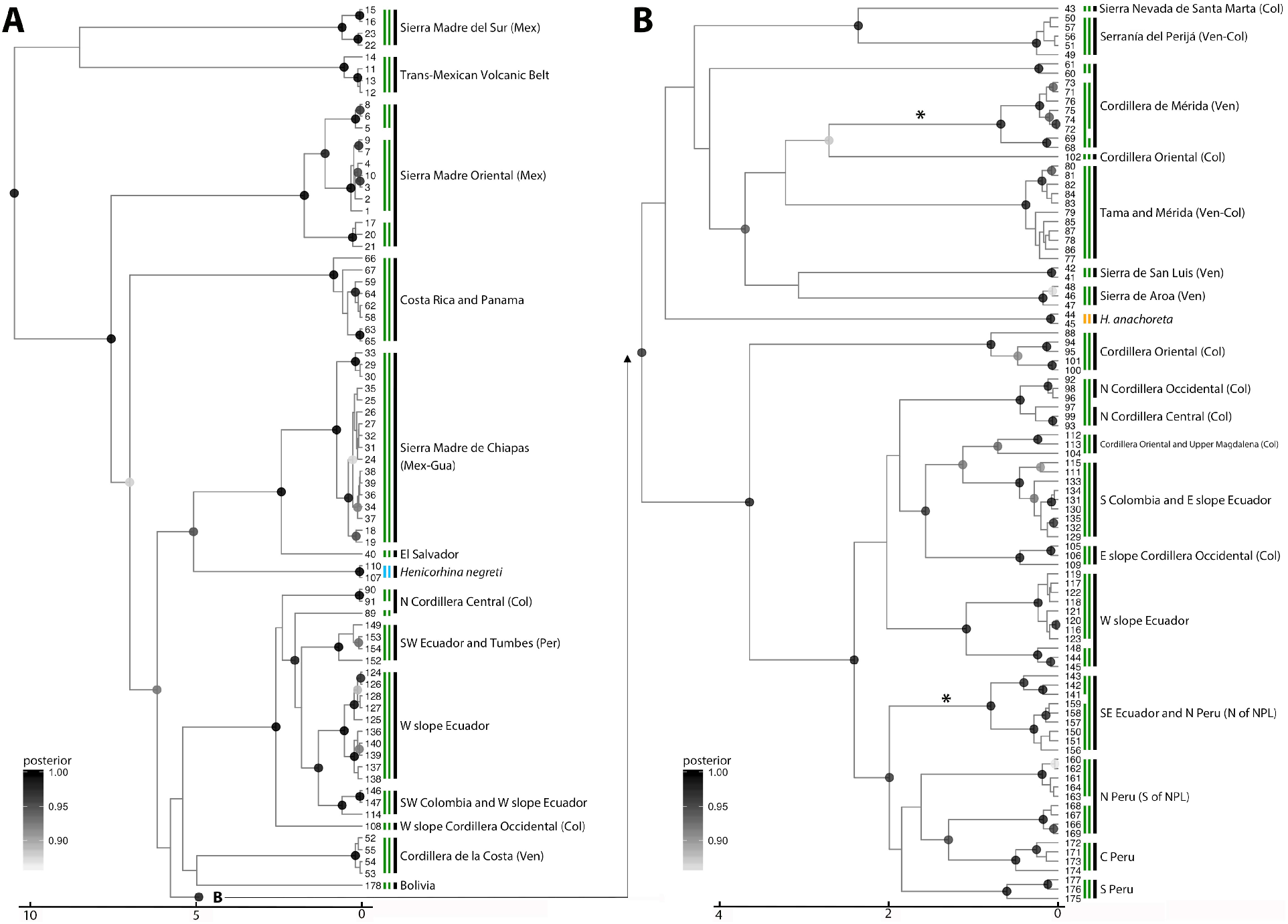
Phylogenetic relationships and divergence times among unique mtDNA haplotypes in the *H. leucophrys* complex inferred using BEAST suggest (1) paraphyly of *H. leucophrys* with respect to *H. anachoreta* and *H. negreti*, (2) a northern origin for the complex with subsequent colonization of South America, and (3) marked population genetic structure partly attributable to geographic isolation mediated by physical barriers. Color shading on nodes corresponds to posterior probabilities ≥ 0.85. Black vertical lines indicate geographic regions; note that all deep branches correspond to clades from mountain regions in Mexico and Central America and that South American populations are also strongly structured. Green vertical lines signal presumptive species identified using multi-rate Poisson Tree Processes (mPTP, left) and the Bayesian General Mixed Yule Coalescent Model (bGMYC, right); results of these analyses were almost identical, with only minor discrepancies in two clades marked with asterisks. Haplotypes are numbered at the tips of the tree; information on specimens having each haplotype is provided in Supplementary Table 1.

Basal relationships among major South American lineages were unresolved or poorly supported, yet some patterns in the region are noteworthy. Whereas populations from some isolated montane systems (e.g. the Serranía de Perijá, or the Venezuelan Cordillera de la Costa and Sierra de San Luis) formed distinct clades, this was not the case for the main cordilleras of the northern Andes, resulting in complicated patterns of area relationships. For example, several populations from the western slope of the Andes from Colombia through Ecuador and into northwestern Peru formed a large clade (clade i. in Figure 2), but this clade was not exclusive because it also included some -but not all-populations from the northern sector of the Cordillera Central of Colombia and did not cluster all populations from the western Andes: *H. negreti* and lineages of *H. leucophrys* from the northern and southern sectors of the Cordillera Occidental of Colombia and from western Ecuador occupied different positions in the tree. Likewise, birds from the Cordillera Oriental of the Colombian Andes formed multiple distinct clades seemingly distantly related to each other, and populations from the Central and Southern Andes (i.e. from Bolivia and Peru south of the Marañón Valley or North Peru Low) formed at least two highly divergent clades with differing affinities. Given weak support for deep branches in South America we do not elaborate further on relationships among major biogeographic areas, but do emphasize the complexity of phylogeographic pattern and the strong genetic structure existing over relatively fine spatial scales throughout the continent.

Part of the complexity in phylogeographic pattern related to occurrence of phylogenetically distant groups in the same regions can be understood by examining elevational distributions: lineages known to replace each other along elevational gradients in northern South America are not sister to each other (Figure 3). This was true of taxa occurring in the Ecuadorean Andes (*H. l. hilaris* and nominate *H. l. leucophrys*), the Santa Marta mountains (*H. l. bangsi* and *H. anachoreta*), and in the western slope of the Colombian Andes (*H. l. brunneiceps* and *H. negreti*). Our data further revealed a previously unknown case of cryptic replacement of mtDNA lineages along an elevational gradient in the Venezuelan Andes. The lineage occurring in the Tamá massif near the Colombia-Venezuela border crosses the Táchira depression to the northeast into the Cordillera de Mérida where we found it from c. 1520 m to c. 1920 m. Only a small distance upslope in this range, a different lineage occupied elevations from c. 2100 m to c. 2750 m. Divergence in mtDNA sequences between lineages replacing each other with elevation was substantial, in all cases exceeding 5% uncorrected *p* distances (Figure 3).

**Figure 3.**
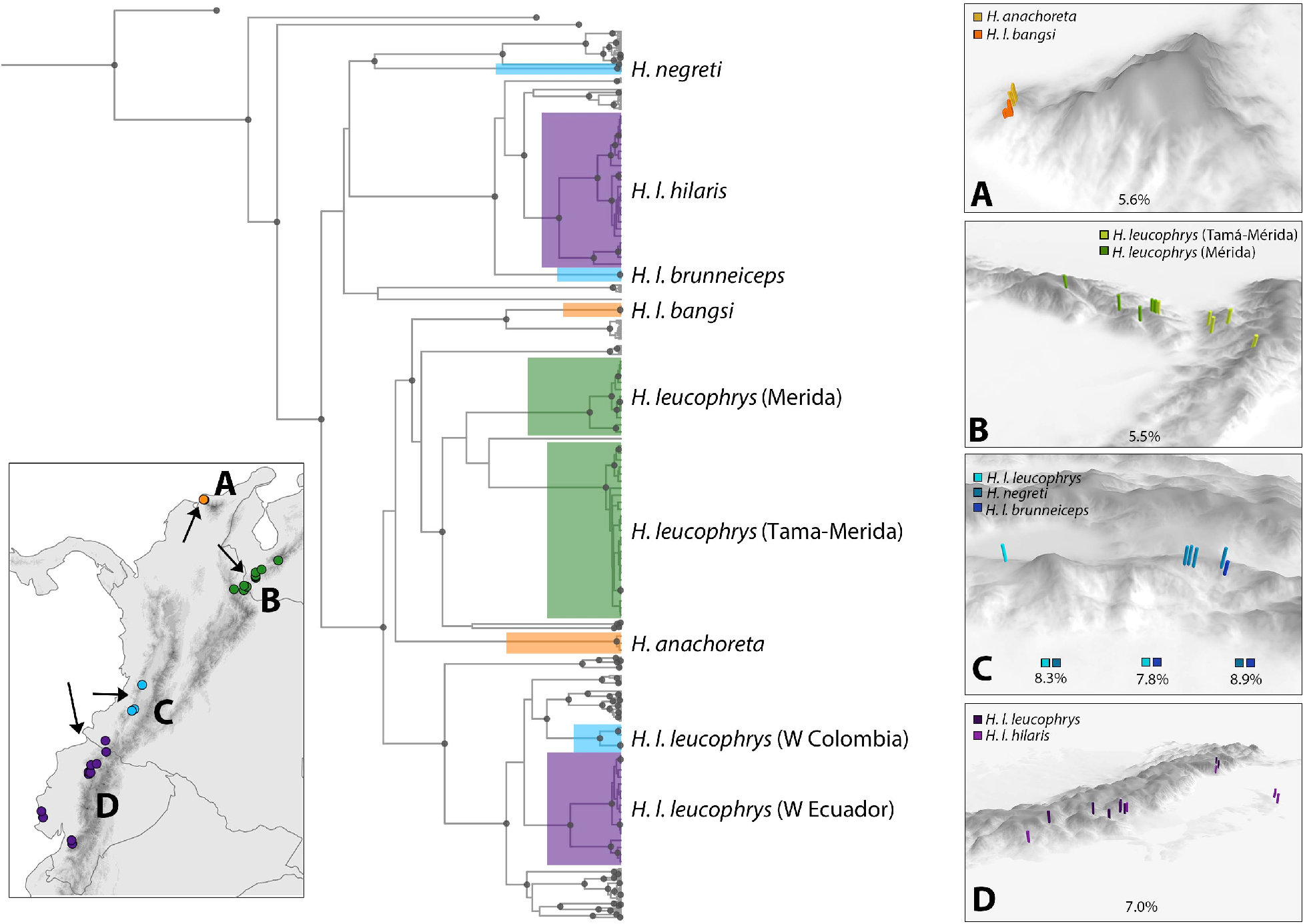
Elevational replacements involving distinct lineages of the *H. leucophrys* complex in montane South America. Lineages replacing each other with elevation in a region share colors in the map, the phylogeny (modified from Supplementary Figure 1, nodes indicated with grey dots have ≥0.85 posterior probability support), and the close-up view of mountain slopes, where different shades are used for each lineage (arrows on the map show the direction from which mountains are seen in panes A-D). In at least three regions (A, C, D), elevational replacements do not involve sister taxa with the only possible exception being the novel case of cryptic replacement of lineages in the Cordillera de Mérida, Venezuela (B), where affinities of lineages to each other and to those from other regions are not strongly supported. Lineages involved in elevational replacements are deeply divergent from each other (panes show mean uncorrected *p* distances in ATPase genes), occur in close proximity, and correspond to different presumptive species identified by coalescent analyses (see text).

In addition to examples of elevational parapatry, our analyses revealed cases where lineages may meet in contact zones along a latitudinal axis. For instance, two lineages differing in c. 7% uncorrected p-distance replace each other along the northern sector of the Cordillera Central of the Colombian Andes. One of these lineages occurs in the northern tip of the cordillera in Antioquia, whereas the other is also found in Antioquia, where we suspect it might range north to the southern extreme of the Aburrá Valley in the outskirts of the city of Medellín. Individuals occupying the northern extreme of the Cordillera Central are most closely allied to geographically distant populations from the western slope of the Andes (i.e. subspecies *hilaris* and *brunneiceps*), which tend to occur at lower elevations and are replaced upslope by nominate *H. leucophrys* in Ecuador or *H. negreti* in southwest Colombia. Several other examples of distinct lineages occurring at different latitudes within mountain systems exist in the Cordillera Oriental of Colombia and along the Andes of Ecuador (Figure 2).

Coalescent approaches to delimit species produced consistent results: both mPTP and bGMYC (the latter with a 0.50 probability threshold to define group membership) recovered *H. anachoreta* and *H. negreti* as distinct species, and both methods identified 37 additional lineages in the *H. leucophrys* complex which may prove to be distinct species (Figure 2, Supplementary Figure 3). Although methods did not exactly agree in how they assigned individuals to presumptive species, congruence was remarkable. The only differences were that in the Cordillera de Mérida, Venezuela, mPTP recognized three presumptive species while bGMYC recognized two, and that in a clade from northern Peru and southeast Ecuador mPTP recognized a single presumptive species and bGMYC recognized two (Figure 2). Applying more stringent probability thresholds to delimit species in bGMYC analyses resulted in the inference of slightly lower numbers of presumptive species: 36 and 35 with 0.90 and 0.95 thresholds, respectively. In general, presumptive species appear to have restricted ranges (Figure 4, Supplementary Figure 4); in some cases, particular mountain systems harbor a single presumptive species (e.g. Sierra Madre del Sur and Trans-Mexican Volcanic Belt in Mexico, Cordillera de la Costa and Sierra de San Luis in Venezuela), but more than one presumptive species may also occur within a region (e.g. Sierra Madre Oriental of Mexico, Sierra Nevada de Santa Marta in Colombia) and a few of them have ranges encompassing various montane areas (e.g. across cordilleras of Costa Rica and Panama). The diversity of presumptive species is especially remarkable in northern South America, with 7-8 identified in Venezuela, 15 in Colombia, and 7-8 in Ecuador. Because our sampling was sparser in Peru and Bolivia, our figures for these countries are likely underestimates of presumptive species richness.

**Figure 4.**
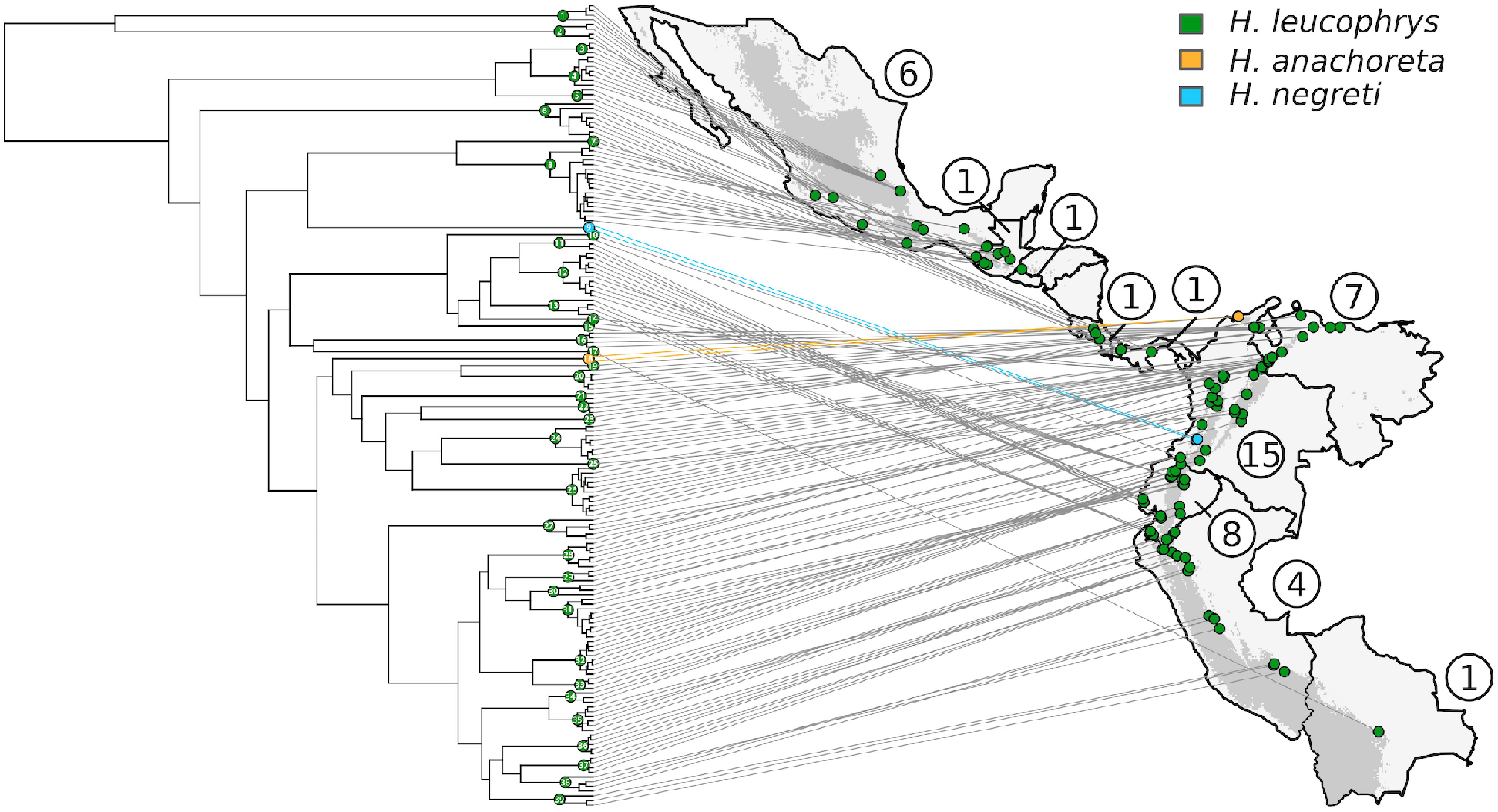
Geographic locations where we sampled 39 presumptive species in the *H. leucophrys* complex identified by coalescent analyses of mtDNA sequences. Dots and numbers on the tree (modified from Figure 2) correspond to species statistically inferred by the Bayesian General Mixed Yule Coalescent Model (bGMYC) with the threshold probability used to define group membership set at 0.50. Colors correspond to species epithets as per the current three-species taxonomy. Encircled numbers on the map indicate the number of presumptive species occurring in each country. Almost identical patterns were observed using the multi-rate Poisson Tree Processes (mPTP) method. For close-up views of geographic locations where each presumptive species was sampled, see Supplementary Figure 4.

Our estimates of divergence times obtained from the BEAST analysis of unique haplotypes indicate that the *H. leucophrys* complex diverged from its sister group (i.e. the clade formed by *H. leucosticta* and *H. leucoptera*) approximately 16.8 million years before present (13.0-20.5 95% highest posterior density, HPD), with the crown age of extant populations dating to 10.6 m.a. (8.4-13.1 95% HPD). Over this period, the complex has diversified into multiple lineages; we found it minimally consists of 10 lineages of at least 5 million years of age and of 26 lineages of at least 2 million years of age (Figure 2, Supplementary Figure 3). The estimated age of the node including all South American populations as well populations from Chiapas, Guatemala, and El Salvador is 6.2 m.a. (5.2-7.3 95% HPD), whereas that of the node including all South American populations excluding *H. negreti* is 5.8 m.a. (4.9-6.7 95% HPD).

Analyses of lineage accumulation over time based on presumptive species identified by bGMYC suggested that rates of diversification in the *H. leucophrys* complex may have declined over time, with a significantly negative gamma statistic (Figure 5). However, through much of the history of the complex, diversification appears to have been nearly constant and exponential, with an apparent downturn in the last million years most likely reflecting that our species delimitation analyses recognized no species younger than this age.

**Figure 5.**
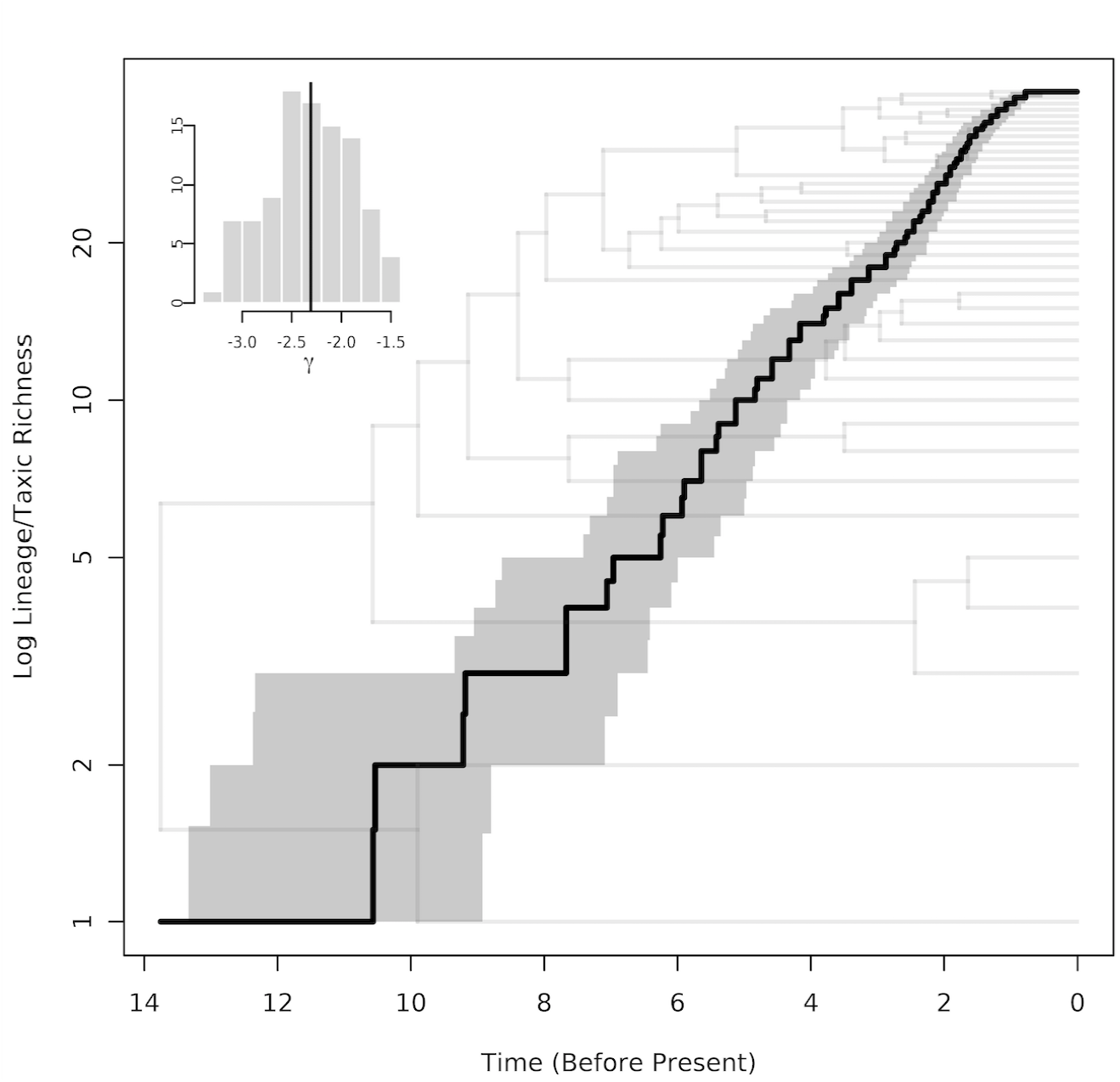
Lineage-through time plot showing accumulation of lineages over time in the *H. leucophrys* complex. Black lines are estimates based on the maximum clade credibility tree and grey indicates the 95% credibility interval across 100 trees for the plot and for estimates of the gamma statistic. The shape of the curve and the associated gamma statistic suggests that rates of lineage accumulation have declined over time, but note that because this analyses used results of bGMYC as input no presumptive species younger than 1 million years were considered. The pattern may also reflect incomplete sampling of young lineages particularly within South America.

## Discussion

### Phylogeography: bridges, barriers and the distribution of genetic variation

As evidenced by the branching pattern in the gene tree with deep splits involving populations from the north, the *H. leucophrys* complex likely originated in the Mexican highlands ca. 8 to 13 m.a., from where it expanded south through Central America, then colonizing South America. Other birds ranging broadly in montane forests also originated in the northern Neotropics, including single species as well as clades which diversified in Central and South America (Pérez-Emán, 2005; Cadena *et al*., 2007; Weir *et al*., 2008; Sánchez-González *et al*., 2015). Molecular-based estimates of when did birds colonize South America from the north vary (Bacon *et al*., 2015; Barker *et al*., 2015); our results indicate colonization by the *H. leucophrys* complex occurred slightly earlier (ca. 7 to 5 m.a.) than a pulse of avian interchange via the Isthmus of Panama 4 to 2 m.a. (Smith & Klicka, 2010). Our estimates of the age of the *H. leucophrys* complex and of the timing of events like its colonization of South America are old relative to what one would expect given published estimates of divergence times among wren genera (Barker, 2017). However, such estimates were derived assuming that *Certhia* and *Troglodytes* diverged ca. 16 m.a. (Moyle *et al*., 2016), while analyses integrating more extensive fossil evidence suggest such divergence occurred much earlier, ca. 27 m.a. (Claramunt & Cracraft, 2015). The time frame for wren diversification implied by the latter analysis is more congruent with our estimated ages for nodes in *Henicorhina* and with previous work in other wren genera based on mtDNA data (Barker, 2007).

Poorly supported relationships among South American clades associated with short internodes subtending long branches are common to *H. leucophrys* and other birds colonizing South America from the north (e.g., Pérez-Emán, 2005; Cadena *et al*., 2007). This indicates range expansions and ensuing rapid diversification of lineages in geographic isolation, a pattern also documented in montane clades with South American (Chaves *et al*., 2011) or uncertain geographic origins (Gutiérrez-Pinto *et al*., 2012). Rapid range expansions occurring in concert across birds may indicate that geological changes like closure of the Isthmus of Panama and uplift of mountains increased connectivity among formerly isolated regions, enabling subsequent diversification of various taxa over the vast South American landscape; climatic changes driving population isolation likely facilitated such diversification (Barrantes, 2009; Ramírez-Barahona & Eguiarte, 2013).

Genetic divergence associated with landscape features isolating montane habitats is another pattern shared by the *H. leucophrys* complex and co-distributed clades (Weir, 2009). Such features include lowland areas in Central America (Cadena *et al*., 2007; Barber & Klicka, 2010), inter-Andean valleys like the Magdalena and Marañón (Gutiérrez-Pinto *et al*., 2012; Benham *et al*., 2015), and alpine areas separating slopes of cordilleras (Parra *et al*., 2009; Valderrama *et al*., 2014). For many Neotropical montane birds that have been studied, genetic structure across geographic barriers coincides with plumage differences (Cadena *et al*., 2011; Winger & Bates, 2015; Winger, 2017). Phenotypic differences among distinct lineages of *H. leucophrys*, however, are either subtle or appear to be nonexistent (Kroodsma & Brewer, 2005). Reduced gene flow across barriers may have influenced vocal differentiation of wood-wrens more strongly, but given their complex songs, confirming it awaits studies documenting repertoires of individuals as well as variation within and among populations. Such data are relevant given uncertainty about species limits in the complex (see below) because vocalizations likely play a critical role in species recognition (Caro *et al*., 2013; but see Halfwerk *et al*., 2016).

In sum, our study and other phylogeographic analyses point to geological and climatic dynamics of the montane Neotropics as drivers of avian speciation both by (1) promoting dispersal across formerly isolated areas and (2) spurring diversification linked to the origin of new habitats resulting from uplift processes and vicariance.

Furthermore, because wood-wrens live in rugged landscapes and disperse little, their populations may become isolated and diverge even without marked geological or climatic changes (Smith *et al*., 2014). Beyond patterns common to the *H. leucophrys* complex and other tropical montane birds, two aspects appear unique to our study system. First, the degree of genetic structure within a single recognized species we uncovered far exceeds that observed in other montane birds. Second, our finding that distinct mtDNA lineages which likely diverged in allopatry have come into contact in various regions and some coexist segregated by elevation is novel. Because the extreme genetic structure we uncovered may imply that *H. leucophrys* comprises more species than traditionally thought and because secondary sympatry of divergent populations is crucial to the buildup of species richness, these results have implications for understanding tropical diversity and the historical and evolutionary processes generating and sustaining it.

### Extreme population structure, cryptic divergence, and patterns in tropical diversity

We uncovered genetic structure in the *H. leucophrys* complex across well-known geographic barriers (Hazzi *et al*., 2018), but also over fine scales in ways not associated with divergence in other tropical montane birds. For example, in the Cordillera Occidental and Cordillera Central of Colombia, where other birds show little to no population structure (Cadena *et al*., 2007; Gutiérrez-Pinto *et al*., 2012; Isler *et al*., 2012; Valderrama *et al*., 2014), we found six mtDNA lineages of at least 1 million years of age. These lineages and others have restricted ranges, and some of their boundaries reflect topographic or climatic breaks (Graham *et al*., 2010; Supplementary Figure 4). Traits affecting dispersal abilities and dependence on closed understory habitats mediate divergence across putative barriers and thus diversification in topographically complex landscapes (Burney & Brumfield, 2009; Smith *et al*., 2014). Because wood-wrens are small-bodied, have small and rounded wings and live in dark forest understory, they likely disperse little (Moore *et al*., 2008), and this may account in part for their exceptionally strong population structure (Claramunt *et al*., 2012; Salisbury *et al*., 2012; but see Smith *et al*., 2017). Deep phylogeographic structure also exists in other small-bodied wrens (i.e. other *Henicorhina, Cistothorus, Troglodytes;* Dingle *et al*., 2006; Campagna *et al*., 2012; Galen & Witt, 2014; Robbins & Nyári, 2014), suggesting that their biology predisposes populations to become isolated and diverge.

Regardless of the ultimate causes of population structure, we discovered heretofore underappreciated diversity within a taxon traditionally treated as a single species. Although our study employed only one molecular marker, some of the lineages we recovered coexist as distinct phenotypic entities exhibiting behavioral barriers to hybridization (Salaman *et al*., 2003; Caro *et al*., 2013; Burbidge *et al*., 2015), implying that several species are involved. Genetic distances (i.e., divergence times) are not appropriate surrogates for reproductive isolation (Roux *et al*., 2016), but we note that wood-wren lineages demonstrating barriers to gene flow in sympatry (i.e. *anachoreta* and *l. bangsi; negreti* and l. *brunneiceps; negreti* and nominate *leucophrys*) last shared ancestors more recently than many other lineages in the complex. It is also remarkable that the phylogeography of *H. leucophrys* resembles that of *Atlapetes* brushfinches (Emberizidae), which also have a northern origin and montane distribution through the Neotropics, and which have diverged into numerous lineages upon colonizing South America (Sánchez-González *et al*., 2015; J. L. Pérez-Emán, unpubl. data). In contrast to *H. leucophrys, Atlapetes* diversified extensively in plumage and this has arguably influenced taxonomy, with researchers recognizing 28 species in the group (Remsen *et al*., 2018). Just as lineages of the *H. leucophrys* complex replace each other in space, species of *Atlapetes* are for the most part allopatric or replace each other sharply along elevational or latitudinal axes in the Andes (Remsen & Graves, 1995), with their ranges often matching those of lineages of *H. leucophrys* uncovered by our study. This comparison serves to illustrate what might be a more general situation in which clades with roughly similar ages and genetic structure (South American *Atlapetes* are actually younger than South American *H. leucophrys*) may be split to different degrees by taxonomists because of differences among clades in the traits birds employ for signaling and in the lability of such traits. In other words, birds like wood-wrens may be under split owing to their conserved plumage and because the role of vocalizations in species recognition remains understudied (see also D’Horta *et al*., 2013).

Our coalescent analyses indeed suggest that taxonomy underestimates species diversity in the *H. leucophrys* complex: we consistently identified 39 presumptive species across methods. Even the 35 presumptive species identified by bGMYC using a more conservative probability threshold of 0.95 represents a quite remarkable figure relative to similar studies conducted in the Neotropics. In 27 clades of lowland birds ranging from Central America across the Andes through much of Amazonia and even into the Atlantic Forest (Smith *et al*., 2014), the mean number of species identified using bGMYC also with a 0.95 threshold was 5.3 (range 1-18); figures were slightly higher in understory birds (11 clades; mean = 6.6, range 3-11 presumptive species) but still much lower than our estimates for the *H. leucophrys* complex. Moreover, in 173 taxonomic species of birds from the New World subject to phylogeographic analyses employing mtDNA data, the largest number of presumptive species identified by bGMYC with a 0.90 threshold was 23 (Harvey *et al*., 2017b; Smith *et al*., 2017), highlighting the *H. leucophrys* complex as a distinct outlier (Figure 6). This is despite our sparse sampling in the Peruvian and Bolivian Andes, where one would expect more lineages exist. We do not argue that all lineages we uncovered are species given existing evidence, but several are candidates for studies examining other molecular markers, morphology, voices, and behavior (Caro *et al*., 2013; Burbidge *et al*., 2015; Halfwerk *et al*., 2016). The allopatric ranges of most wood-wren lineages preclude tests of intrinsic barriers to gene flow, but given postzygotic isolation in other phenotypically cryptic, old lineages of Neotropical birds (Pulido-Santacruz *et al*., 2018), some of them may well be reproductively isolated.

**Figure 6.**
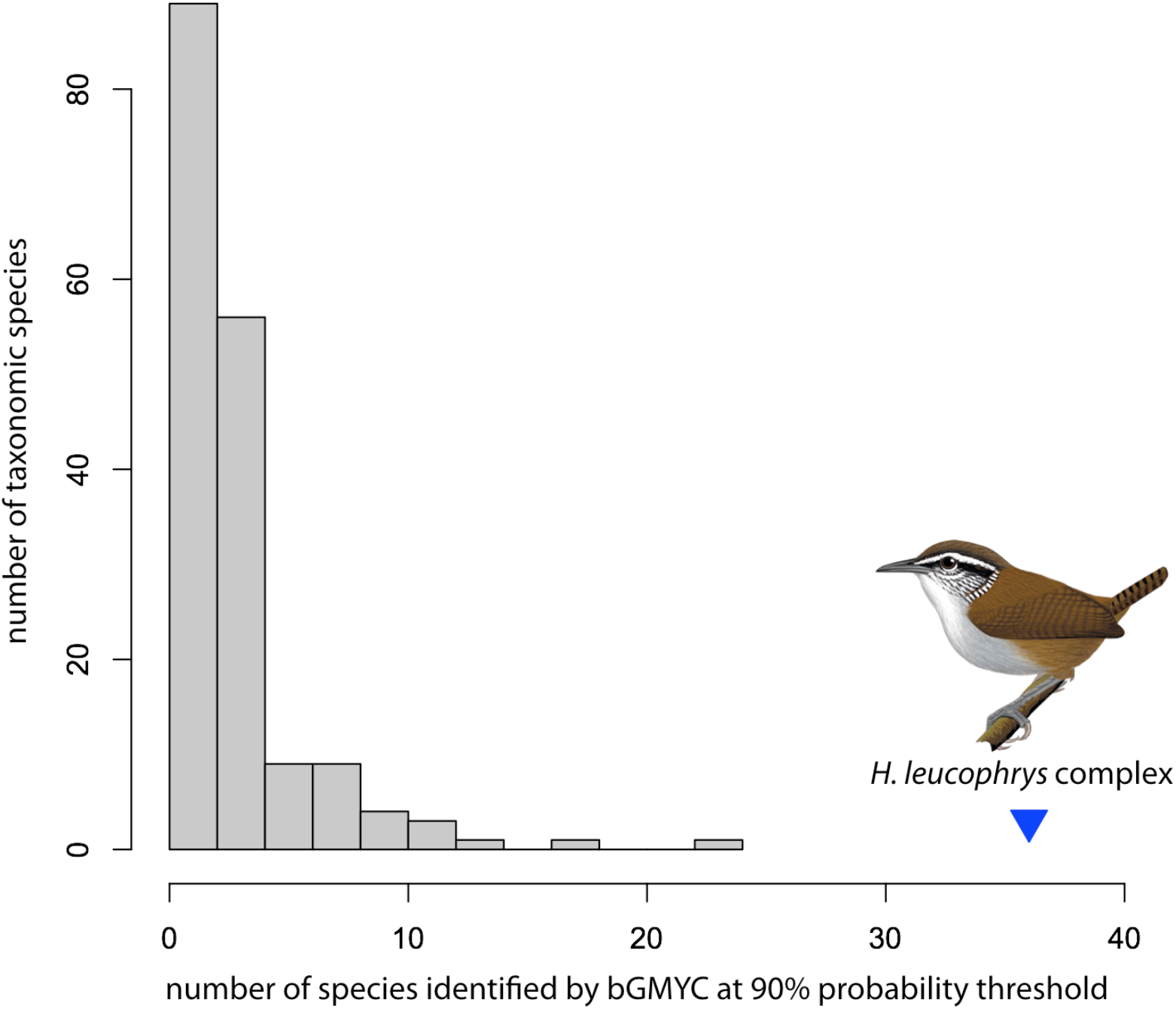
Relative to the frequency distribution of the number of presumptive species identified by coalescent analyses of mtDNA sequence data within 173 taxonomic species of New World birds (data from Harvey et al. 2017, Smith et al. 2017), our result for the *H. leucophrys* complex is a distant outlier. Even if many lineages cannot be shown to be reproductively isolated from others, the data reveal a remarkable and previously undocumented degree of population genetic structuring. Illustration by F. Ayerbe.

Even if hypothetical species our analyses flagged represent distinct lineages not yet reaching the status of “biological” species (Carstens *et al*., 2013; Sukumaran & Knowles, 2017), our work allows for conclusions about cryptic diversity which should be robust to analyses involving other data. First, regardless of the species concept one follows and of the criteria one uses to recognize species, there are more wood-wren species than traditionally thought. Second, the *H. leucophrys* complex comprises multiple independently evolving populations which have diverged to different degrees along the speciation continuum; if one embraces the generalized lineage concept, which views species as segments of metapopulation lineages and considers all other species “concepts” as contingent –albeit not necessary– properties of species one may use as criteria to recognize them (de Queiroz, 1998, 2007), then the complex is arguably a collection of several kinds of species, all of which represent fundamental evolutionary units. Finally, even if one takes a conservative standpoint and treats most of the lineages we uncovered as distinct populations of a single or a few species (Sukumaran & Knowles, 2017), the *H. leucophrys* complex has clearly differentiated into numerous lineages (especially following its colonization of South America) and such lineages have persisted over long periods.

Because most wood-wren lineages are not sympatric (see below), our finding that the *H. leucophrys* complex probably comprises multiple species has no influence on local estimates of diversity. However, species richness and endemism at regional scales might need to be revised. For example, current taxonomy recognizes only one species (and only five subspecies) in the complex in Venezuela (Kroodsma & Brewer, 2005), yet we uncovered 7-8 presumptive species –each endemic to a particular montane system– in the country. If lineages identified as presumptive species are elevated to species status and similar patterns exist in other taxa not yet studied, then geographic variation in population structure may alter knowledge of spatial patterns of diversity (e.g., differences in species richness among cordilleras and slopes of cordilleras of Colombia; Kattan *et al*., 2004; Figure 4, Supplementary Figure 4). Alternative classifications also affect inferences about beta diversity; under current taxonomy, there is no species turnover along thousands of kilometers and across multiple geographic barriers from Mexico to Bolivia except for the local replacements involving *H. anachoreta* and *H. negreti*. At the other extreme, if distinct lineages of *H. leucophrys* are species, then spatial turnover would be substantial even over relatively short distances (e.g., in Colombia), likely exacerbating differences among regions in beta diversity (Gaston *et al*., 2007; Fjeldså *et al*., 2012) and with potential conservation implications (Socolar *et al*., 2016). Recognizing distinct lineages as species would also alter the perceived role of features of montane landscapes setting range limits and thus explaining spatial turnover in assemblages (Graham *et al*., 2010).

In addition to affecting perceptions of patterns of diversity, our results have implications for thinking about historical processes underlying such patterns. The accumulation of biological diversity via diversification within a region like the montane Neotropics requires that (1) populations become isolated to initiate divergence, (2) budding population isolates persist in time, (3) populations expand their ranges and come into secondary sympatry, and (4) newly sympatric populations are differentiated enough that they may coexist without coalescing owing to hybridization or without excluding each other via competition (Mayr, 1942; Ricklefs & Bermingham, 2007). In the following we discuss our results in the context of these steps.

### Lineage splitting, persistence, and the origin of diversity

A leading explanation for high tropical diversity involves latitudinal differences in net diversification rates (Fischer, 1960; Schluter & Pennell, 2017). In particular, rapid diversification may explain the high species richness and concentration of narrow-ranged species of birds in tropical mountains, which cannot be accounted for by area or contemporary climate (Jetz *et al*., 2004; Fjeldså *et al*., 2012). Although evidence that diversification rates vary with latitude remains mixed in birds (Ricklefs, 2006; Martin & Tewksbury, 2008; Jetz *et al*., 2012; Belmaker & Jetz, 2015) and other taxa (e.g. Pyron & Wiens, 2013; Pyron, 2014; Rolland *et al*., 2014; Schluter, 2016; Rabosky *et al*., 2018), tropical mountains are indeed hotbeds of rapid diversification (Madriñán *et al*., 2013). Furthermore, differences in the rate at which species originate may not be as important as the rate at which they go extinct in establishing broad-scale patterns in avian diversity (Hawkins *et al*., 2006; Weir & Schluter, 2007; Pulido-Santacruz & Weir, 2016). Accordingly, the high diversity and endemism of tropical montane areas may reflect low extinction rates of species (Fjeldså *et al*., 2012). In turn, higher diversification rates at higher elevations in montane areas (Quintero & Jetz, 2018) may reflect both high speciation and low extinction (Fjeldså & Irestedt, 2009). A complementary historical explanation for diversity in tropical mountains which is less commonly addressed in the literature is high persistence of budding populations, an important control of rates of speciation (Mayr, 1963; Dynesius & Jansson, 2014; Rabosky, 2016).

We found that the *H. leucophrys* complex radiated rapidly into multiple lineages, several of which have persisted for periods exceeding millions of years. Also, LTT plots suggest nearly constant rates of exponential accumulation of lineages over nearly 10 million years, with an apparent slowdown in diversification in the last million years. Although LTT plots with such a shape and their associated negative gamma statistic are often considered evidence of ecological limits to diversification (Rabosky & Hurlbert, 2015), we interpret the pattern more as an artifact of our methods resulting from (1) using species based on a model specifying a divergence threshold separating population-level processes (gene coalescence) from diversification dynamics (speciation and extinction) as units for analysis (Reid & Carstens, 2012), and (2) potential limitations in geographic sampling leading to failure to identify additional independent lineages of young age. Because the bGMYC analysis we employed to delimit presumptive species established an age cutoff of ca. 1 m.a. defining the units included in the LTT analysis, we simply conclude that diversification was nearly constant through much of the history of the *H. leucophrys* complex. To the extent that similar diversification dynamics may characterize evolutionary history of other Neotropical montane birds, high rates of lineage splitting (Harvey *et al*., 2017b) and high persistence of such lineages over time (Smith *et al*., 2017) have likely contributed to diversification and probably account for the high diversity of tropical montane systems and, more broadly, to large-scale biodiversity patterns such as latitudinal gradients in species richness.

### Range dynamics, secondary sympatry and the regional buildup of diversity

Our data revealed that mtDNA lineages in the *H. leucophrys*, which likely diverged in geographic isolation, have come into secondary sympatry. This is most evident where divergent mtDNA lineages not sister to each other segregate with elevation. In addition to previously documented cases of elevational replacements of lineages involving distinct taxa (i.e. different species or subspecies in the Sierra Nevada de Santa Marta, in western Colombia, and in western Ecuador), we discovered a novel elevational replacement of distinct lineages in the Mérida Cordillera of Venezuela where no phenotypic differences had been noted. Likewise, previous work in other wood-wrens revealed that although *H. leucoptera* is nested within *H. leucosticta*, the lineage of *H. leucosticta* replaced by *H. leucoptera* at higher elevations in the Cordillera del Cóndor east of the Andes is distantly related to it, whereas its closest relative seemingly occurs in the Chocó region west of the Andes (Dingle *et al*., 2006). The consistent pattern of elevational replacements involving fairly distant relatives as opposed to sister lineages fits the hypothesis that evolutionary divergence in tropical montane birds occurs largely in allopatry and not in parapatry along mountain slopes (Patton & Smith, 1992; García-Moreno & Fjeldså, 2000; Caro *et al*., 2013).

In addition to documenting elevational replacements, we found evidence of regional co-occurrence of lineages replacing each other with latitude (e.g. along the cordilleras of Colombia and Ecuador). More fine-scaled sampling is required to determine whether geographic gaps separate the ranges of such lineages or if they come into close contact. Part of the observed genetic differentiation along the latitudinal axis may reflect the propensity of the linear distributions of tropical montane birds to become fragmented (Graves, 1988). However, some lineages replacing each other with latitude in a region are not sisters and may even be distantly related, which suggests range expansions and secondary contact rather than primary divergence along cordilleras.

Other intriguing phylogeographic patterns aside from secondary contact of lineages in elevational or latitudinal parapatry speak to the dynamism of geographic ranges over broad scales. For example, we found that populations of H. *leucophrys* from southern Mexico (Chiapas), Guatemala and El Salvador are not closely related to other Middle American populations; within a large, otherwise South American clade, these specimens appeared closest to *H. negreti*, a species endemic to western Colombia whose northernmost records are ca. 1700 km south of montane El Salvador. Likewise, the only sequence analyzed from Bolivia is a long branch more closely allied to lineages from northern South America (Colombia and Venezuela) than to geographically much closer lineages from Peru. Because closest relatives may occur in distant areas, spatial patterns of genetic variation are not easily accounted for by geography (e.g. by isolation-by-distance; Seeholzer & Brumfield, 2018). Given that such patterns are unlikely evidence of long-distance dispersal and are not unique to wood-wrens in the region (Cadena *et al*., 2007), considering dynamics of expansion and contraction of geographic ranges involving localized extinctions is crucial to understand biogeographic and demographic processes underlying the distribution of genetic and species diversity in Neotropical birds.

Shifting climatic conditions affecting habitat connectivity drive changes in species ranges, thereby influencing phylogeographic patterns and the buildup of montane diversity (Ramírez-Barahona & Eguiarte, 2013; Flantua & Hooghiemstra, 2018). Species ranges may also experience phases of expansion and contraction linked to shifts in ecological specialization and interactions with natural enemies (i.e. the taxon cycle; Wilson, 1959; Ricklefs & Bermingham, 2002). Although taxon cycles are more evident in insular settings with discrete populations and areas (e.g., Ricklefs & Bermingham, 1999; Jønsson *et al*., 2014), they may also take place in continents (Graves, 1982). In fact, lineages experiencing the taxon cycle may account for what one might call continental great speciators like *Henicorhina* wood-wrens, which occur widely in space -revealing an ability to expand their ranges-yet split into isolated populations at a fast rate due to cessation of gene flow (cf. Diamond *et al*., 1976). Wood-wrens disperse little at present because of their morphology and ecology, which arguably explain their remarkable patterns of genetic structure reflecting long-term population isolation. However, our findings that wood-wrens dispersed throughout much of the montane Neotropics from a northern area of origin and that several lineages achieved secondary sympatry indicate that episodes of range expansion interspersed with periods of divergence occurred at various moments, possibly in sync with morphological or behavioral changes influencing their abilities to disperse (Pigot & Tobias, 2015; Hosner *et al*., 2017). Furthermore, gaps separating the ranges of closely related lineages of wood-wrens arguably reflect extinctions of intervening populations of formerly widespread lineages, which left vacant spaces that could, in turn, become occupied by other expanding lineages.

Phylogeographers will often not detect range dynamics embodied in the taxon cycle because incomplete reproductive isolation between young lineages can result in homogenization of gene pools upon secondary contact (Kearns *et al*.,2018). Furthermore, niche similarities between incipient species achieving contact may preclude long-term sympatry owing to interspecific competition (Pigot & Tobias, 2013). Irrespective of whether the patterns we observed resulted from the taxon cycle, we identified aspects making the *H. leucophrys* complex well suited for further work on the origins of tropical diversity and its accumulation over time and space. Our results and other work on the complex reveal that the completion of reproductive isolation between lineages meeting in secondary sympatry seemingly exhibits a continuum ranging from neutral divergence with no obvious phenotypic differences (forms in montane Venezuela), to phenotypic and behavioral divergence with persistent interbreeding (*hilaris* and nominate *leucophrys* in Ecuador), to completed speciation with little to no hybridization (anachoreta and negreti vs. various forms of leucophrys in Colombia; Salaman *et al*., 2003; Dingle *et al*., 2008; Dingle *et al*., 2010; Caro *et al*., 2013; Burbidge *et al*., 2015; Halfwerk *et al*., 2016). Furthermore, divergence in elevational ranges occurring during periods of isolation (Cadena, 2007; Tobias *et al*., 2014) or arising in secondary sympatry (Diamond, 1973; Freeman, 2015) has enabled coexistence of lineages at the landscape scale in various regions. Given that range boundaries may be maintained –and possibly reinforced– evolutionarily by phenotypic and behavioral barriers to interbreeding and ecologically by competition (Jankowski *et al*., 2010), our study has uniquely captured wood-wren populations in the act of building up diversity via divergence and persistence in allopatry, achievement of secondary sympatry, and coexistence mediated by ecological and evolutionary divergence. Comparative work on the structure and dynamics of contact zones between lineages should provide rich insights into the origin and maintenance of high diversity in tropical mountains.

## Acknowledgements

We are grateful to the institutions allowing access to tissue samples for our study: Academy of Natural Sciences of Drexel University, Burke Museum of Natural History and Culture at University of Washington, Colección Ornitológica Phelps, The Field Museum, Instituto Alexander von Humboldt, Instituto de Ciencias Naturales at Universidad Nacional de Colombia, Louisiana State University Museum of Natural Science, Museum of Vertebrate Zoology at UC Berkeley, Peabody Museum of Natural History at Yale University, and the Zoological Museum at the University of Copenhagen. We thank field collectors and field assistants for their crucial role in obtanining samples, as well as researchers who conducted groundbreaking work on our study system, made their sequences available and assisted us in various ways: Caroline Dingle, Lina Caro, and colleagues. Our work received financial support from the Facultad de Ciencias at Universidad de los Andes, and the Centro de Desarrollo Científico y Humanístico de la Universidad Central de Venezuela. Permits were granted by the Oficina Nacional de Diversidad Biológica, Ministerio del Poder Popular para el Ambiente, Venezuela.

---

**Supplementary Figure 1.**
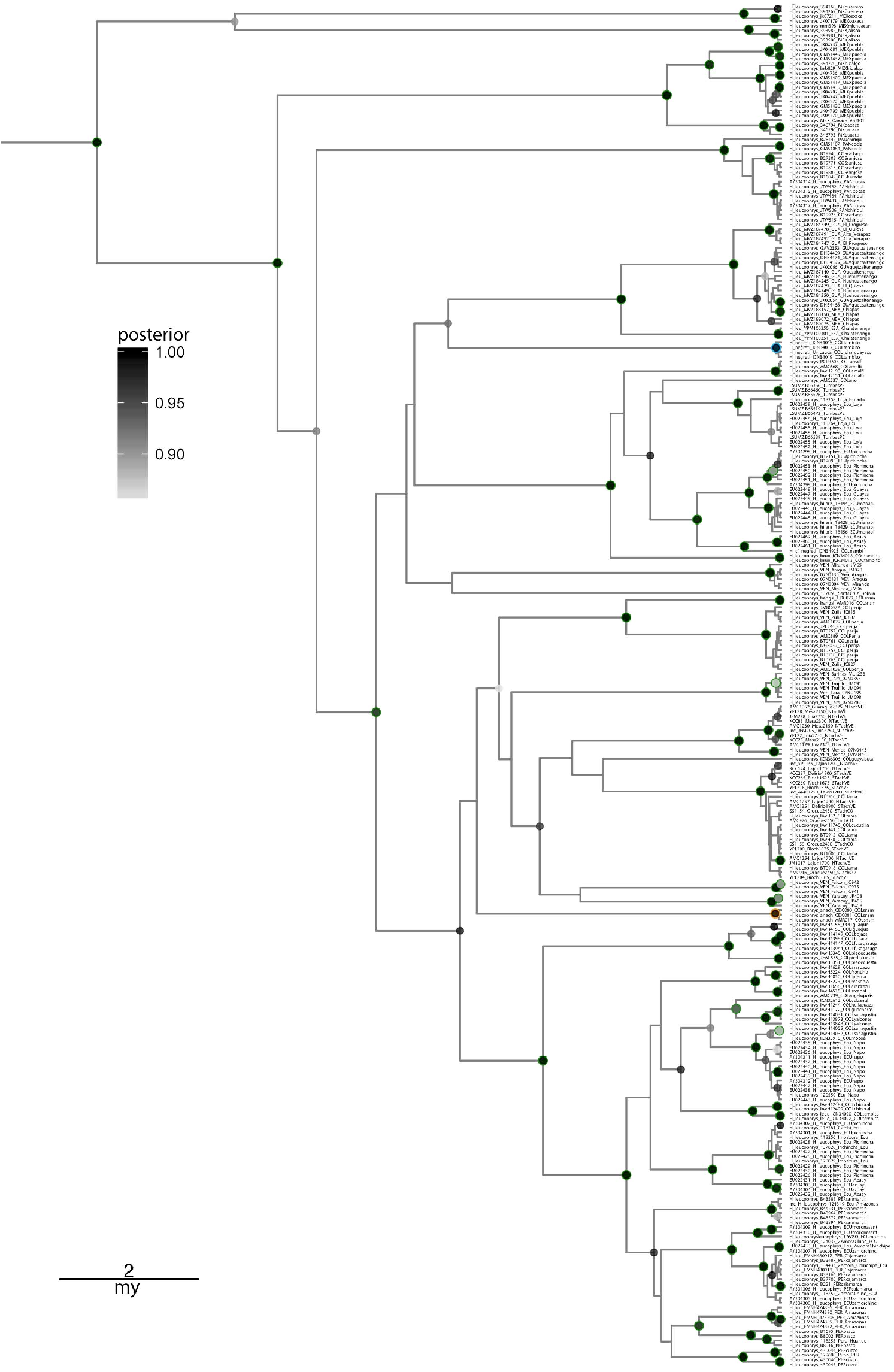
Phylogenetic relationships among individuals in the *H. leucophrys* complex inferred using Bayesian analysis of sequences of the ATPase 6&8 mitochondrial genes. The phylogeny is the maximum clade credibility tree obtained in BEAST. Nodal support (i.e, posterior probabilities ≥ 0.85) is shown using a grey scale. Nodes with a colored outline (green = *H. leucophrys*, blue = *H. negreti*, orange =*H. anachoreta*) were also recovered with strong support (≥ 80% boostrap) in maximum-likelihood analysis (Supplementary Figure 2).

**Supplementary Figure 2.**
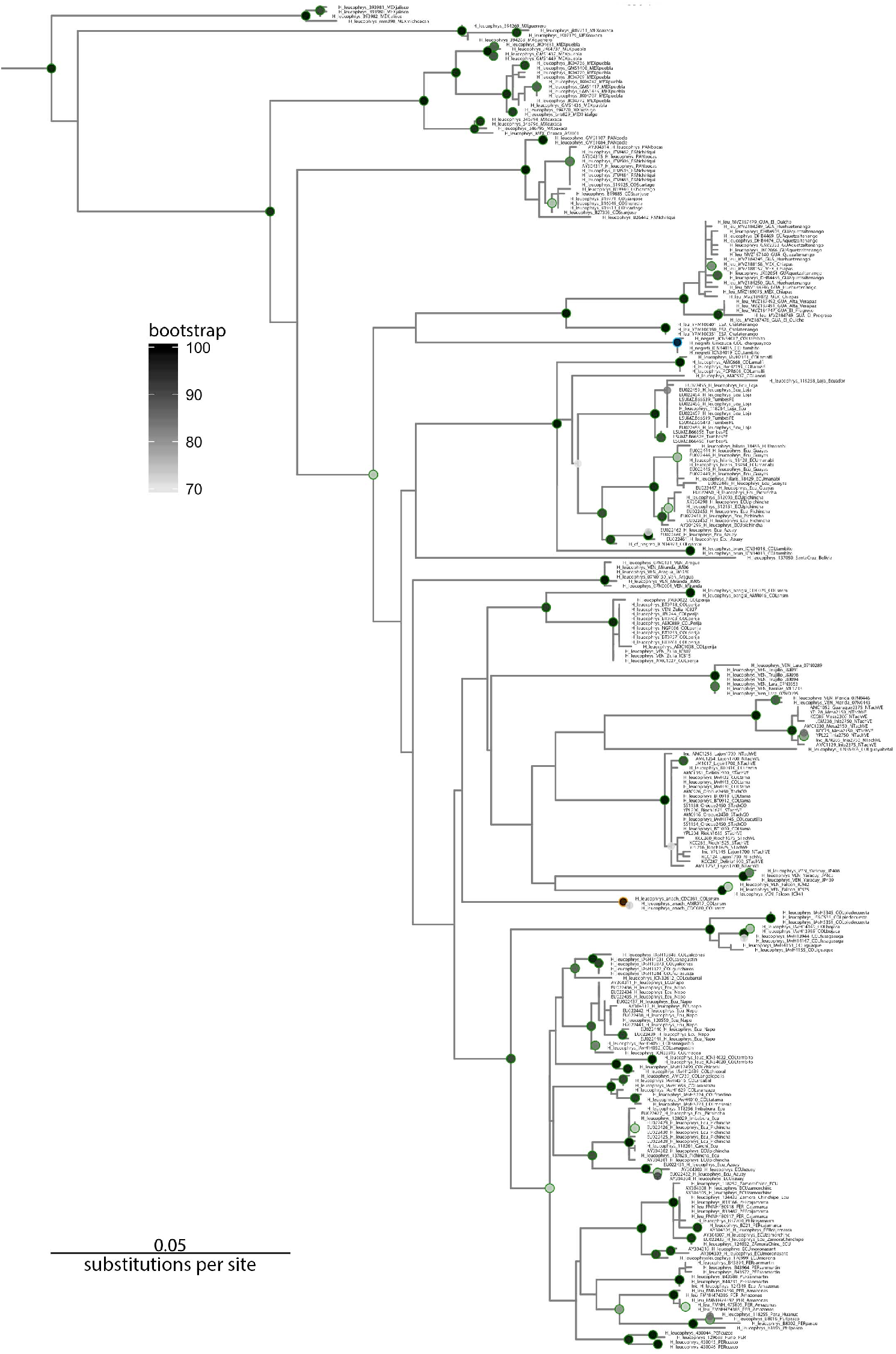
Phylogenetic relationships among individuals in the *H. leucophrys* complex inferred using maximum-likelihood analysis of sequences of the ATPase 6&8 mitochondrial genes. The phylogeny is the maximum-likelihood tree obtained in RAxML. Nodal support (i.e, boostrap values ≥ 80%) is shown using a grey scale. Nodes with a colored outline (green = *H. leucophrys*, blue = *H. negreti*, orange = *H. anachoreta*) were also recovered with strong support (≥ 0.90 posterior probability) in Bayesian analysis (Supplementary Figure 1).

**Supplementary Figure 3.**
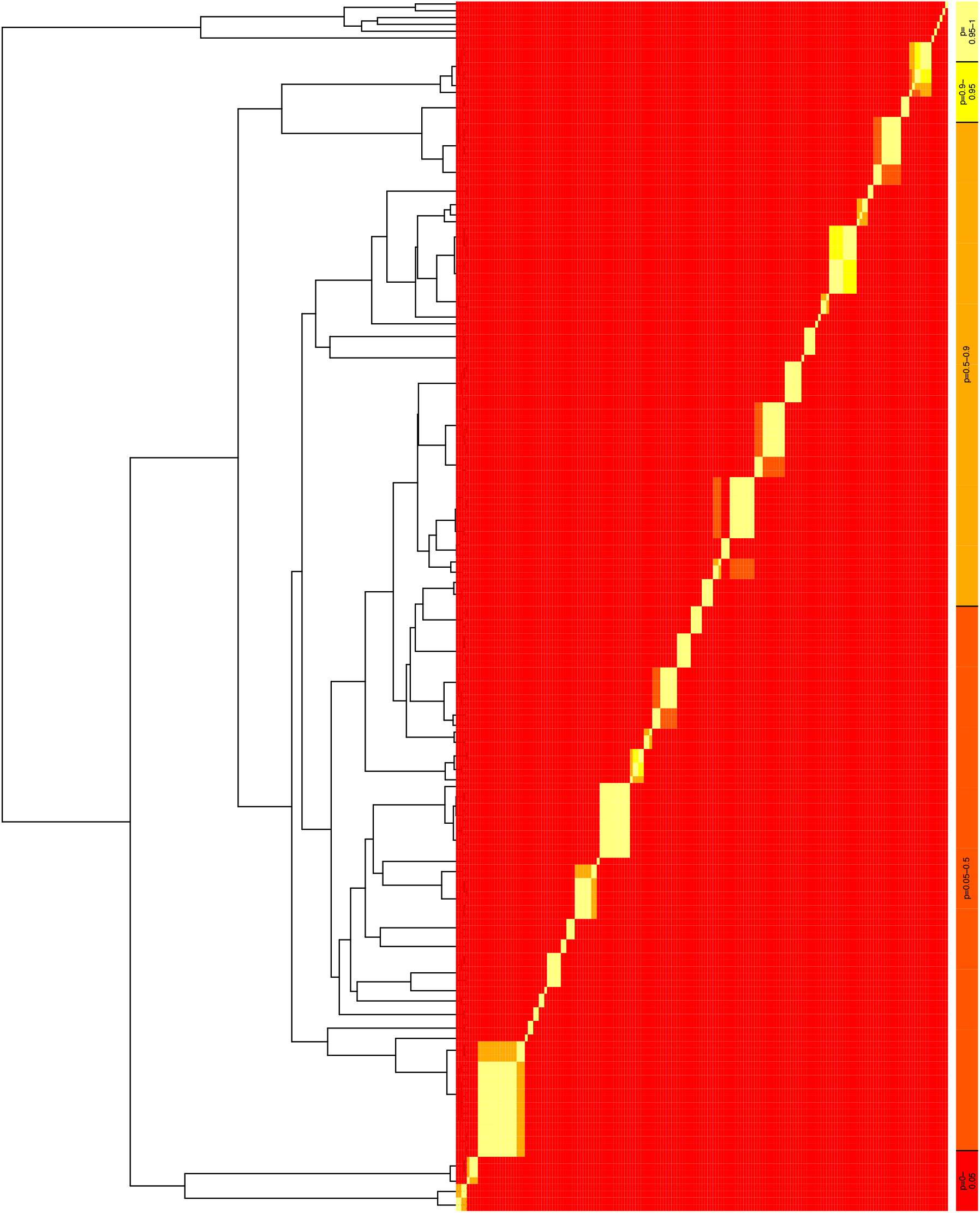
Results of species delimitation analysis in the *H. leucophrys* complex employing the Bayesian General Mixed Yule Coalescent Model (bGMYC). The phylogeny showing relationships among haplotypes is the maximum clade credibility obtained using BEAST and the table to the right is a sequence-by-sequence matrix in which cells are color-coded to indicate the posterior probability that each pair of sequences is conspecific.

**Supplementary Figure 4.**
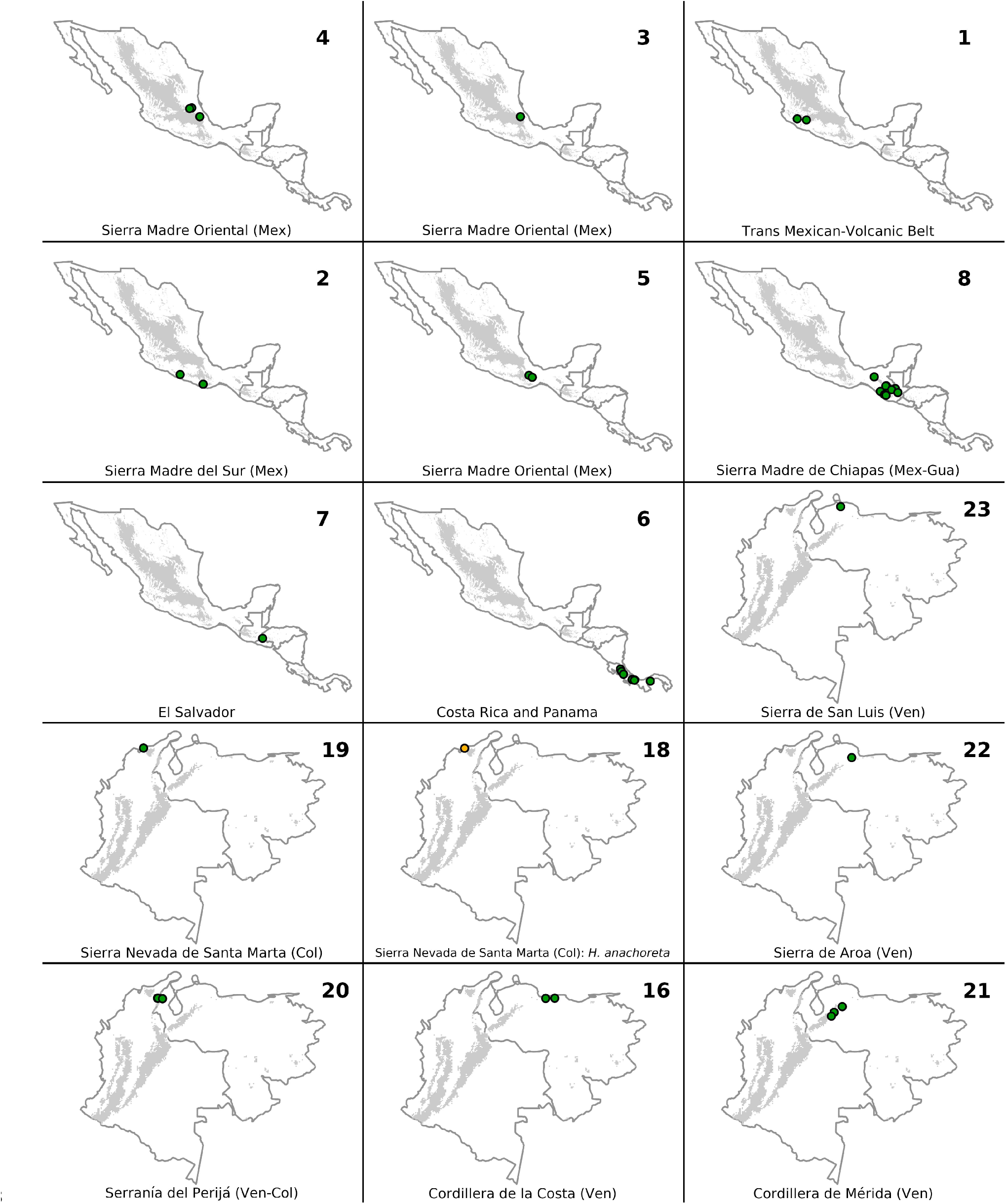
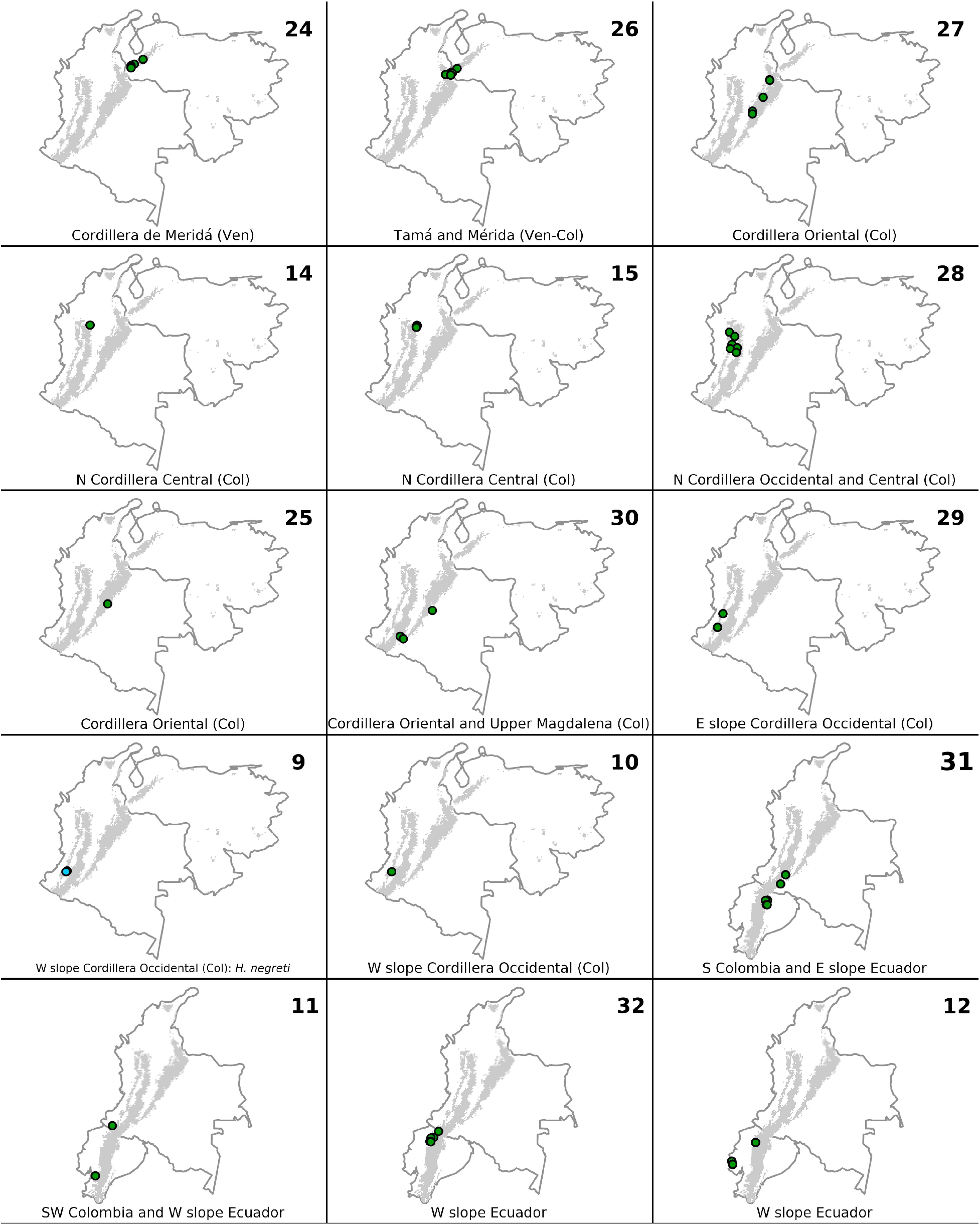
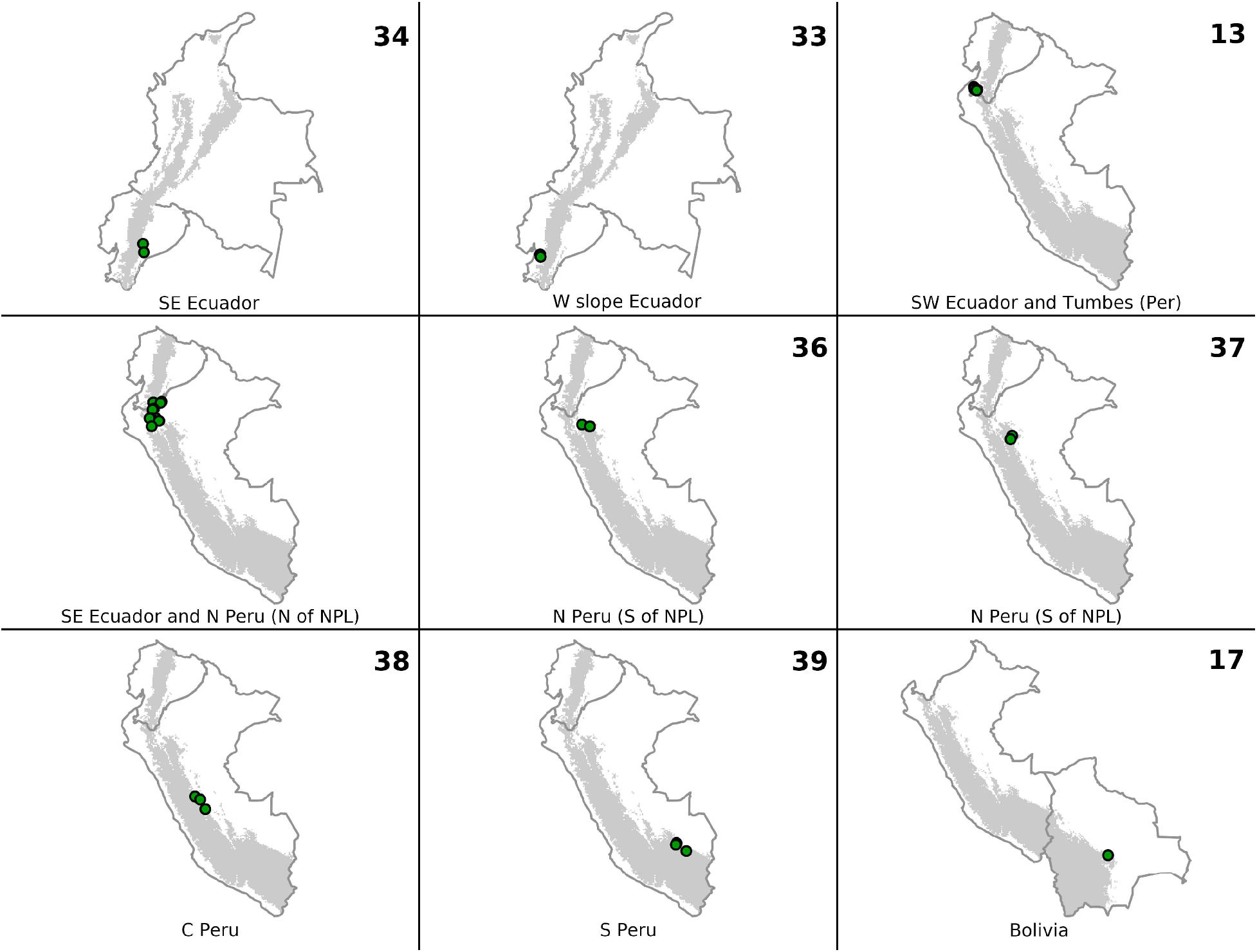
Close-up views of geographic locations where presumptive species in the *H. leucophrys*complex identified by coalescent analyses of mtDNA sequences were sampled. Maps showing known locations of each presumptive species are ordered roughly from North to South, and are numbered according to numbers on nodes in the tree in Figure 4; points on maps are colored based on the current three-species taxonomy recognizing *H. leucophrys, H. negreti*, and *H. anachoreta.* Some presumptive species have relatively large ranges (e.g. no. 6 across Costa Rica and Panama) whereas others appear to be much more restricted, in some cases found at single localities so far (e.g. no. 14 and 15 in the northern extreme of the Cordillera Central in Antioquia, Colombia).

**Supplementary Figure 5.**
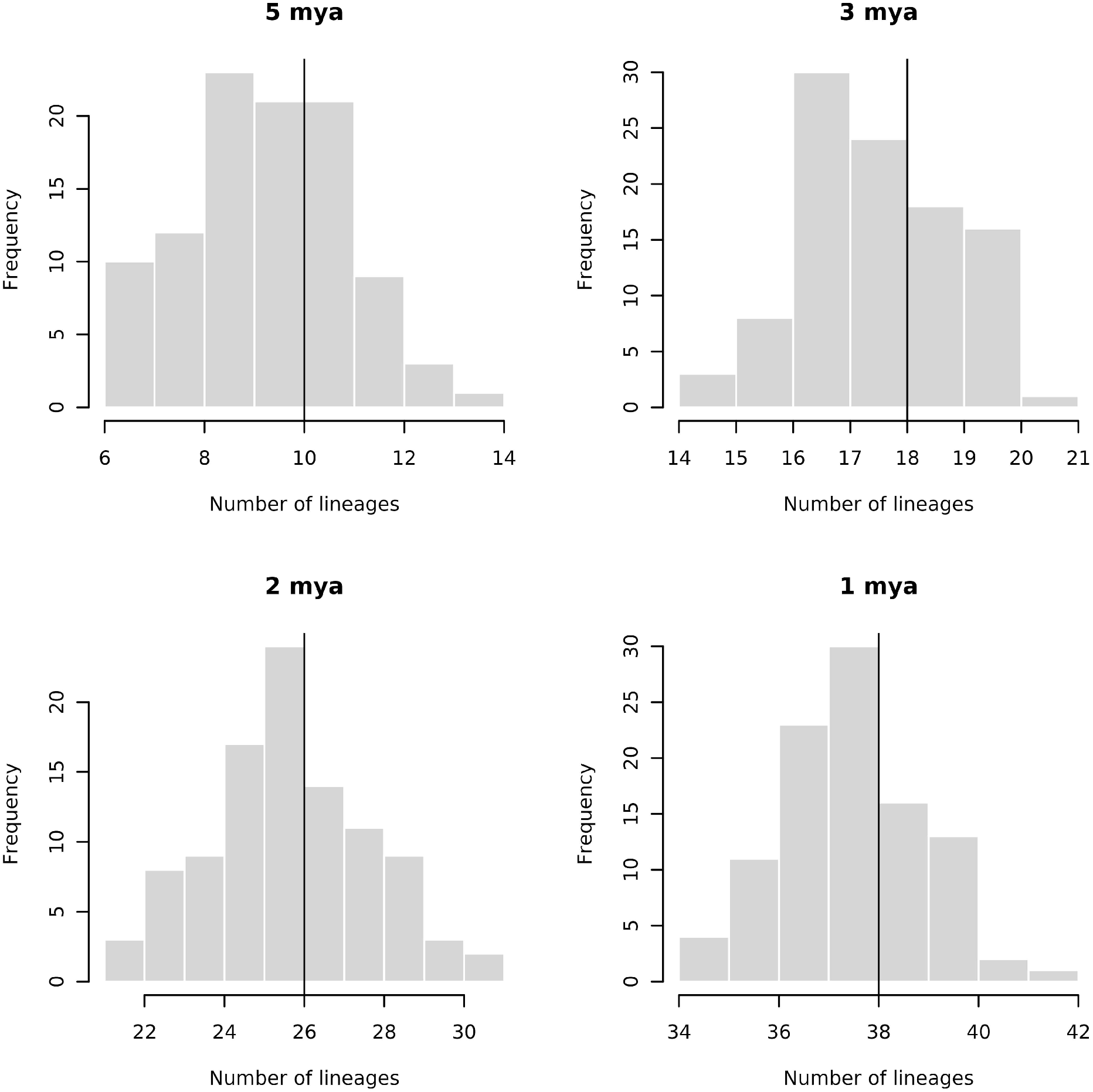
Number of mtDNA lineages in the *H. leucophrys* complex of various ages (from 5 to 1 million years ago [m.a.]). Vertical black lines correspond to the median number of lineages dating to at least each of the four ages (i.e. splitting from their common ancestor with other lineages before each age) observed in a sample of 100 trees in the posterior distribution obtained using BEAST; gray bars are the frequency distributions of number of lineages per age across all trees.

**Supplementary Table 1.** (Provided as a separate .xlsx file). Information on specimens considered in phylogeographic analyses including museum catalogue numbers, locality data, and GenBank accession numbers when available (those for sequences generated for this study are pending). For each specimen, we also indicate the name used to refer to it in Supplementary Figures 1 and 2, the ATPase 6/8 haplotype as shown in Figure 2, and the ID of the presumptive species identified using bGMYC to which it belongs (Figure 3, Supplementary Figure 4).

